# Protein degradation by small tag artificial bacterial E3 ligase

**DOI:** 10.1101/2024.09.07.611797

**Authors:** Zhenyi Liu, Ming-Chi Tsai, Soumitra Ghosh, Jessica Lawrence, Sarah Chu, Baris Bingol, Søren Warming

**Affiliations:** Departments of Molecular Biology, Genentech, South San Francisco, CA 94080; Neuroscience, Genentech, South San Francisco, CA 94080

## Abstract

Targeting of proteins for degradation in a reversible manner is a powerful approach to decipher gene function and mimic drug effects, with great potential for drug target discovery and validation. A generalized approach is to tag a protein of interest and then use this tag to recruit an endogenously or exogenously expressed E3 ligase for its polyubiquitination and subsequent degradation via 26S proteasome. However, the often bulky size of the tag and the great variability of substrate-dependent degradation efficiency of mammalian E3 ligases pose great challenges in practice. Here we show that small tags (10-15 amino acids) can be used to efficiently tag endogenous proteins for degradation when coupled with an exogenously expressed <u>a</u>rtificial <u>b</u>acterial <u>E</u>3 ligase (ABEL) consisting of a tag-interacting moiety and the catalytic domain of the bacterial E3 ligase IpaH9.8. We name this versatile and efficient platform <u>de</u>gradation by <u>s</u>mall tag <u>ABEL</u> (DESTABEL). Furthermore, we show that an ABEL containing a nanobody against human α-synuclein mediates efficient degradation in primary neurons as well as in the adult mouse brain. Taken together, our data show that tag-dependent and independent ABELs are powerful yet flexible tools for studies of protein function and drug target validation.

## INTRODUCTION

Manipulation of protein-coding genes can be achieved at three different levels: DNA, RNA or protein, each with its own advantages and limitations. With most genomic loci having only two copies in normal somatic cells, it is relatively easy to knock out the function of a gene at the genomic DNA level, especially with the advent of CRISPR/Cas9-based technology^1^. However, such genetic manipulations are irreversible, and therefore not amenable to temporal control.

Another major concern with sustained perturbation of gene function is the unintentional activation of compensatory mechanisms that mask the true function of the gene under study^2^. In contrast, genetic manipulation at the mRNA level with siRNA^3,4^, shRNA^5^ and antisense oligonucleotides^6^ offers simple and powerful approaches to modulate gene function in a reversible manner, yet the existence of off-target effects remains an unsolved issue^7^.

Alternatively, gene expression and thus protein level could be regulated by transcriptional silencing (*e.g.* CRISPRi). Nevertheless, manipulations of gene expression share the limitation of RNA interference in the inability to efficiently target proteins with a long half-life.

In recent years, several technologies aiming at modulating gene function at the protein level with targeted protein degradation have been described. Many of these technologies take advantage of the ubiquitin-proteasome system, whose function is to maintain protein homeostasis by degrading damaged, misfolded or short-lived proteins^8^. In general, proteins destined for degradation need to be poly-ubiquitinated before being degraded by the 26S proteasome. The poly-ubiquitination process involves three sequential reactions: first, E1 ubiquitin-activating enzyme catalyzes the formation of a thioester bond between the cysteine at its activation site and the C-terminus of ubiquitin. Second, activated ubiquitin is transferred to the E2 ubiquitin-conjugating enzyme. Finally, an E3 ligase, in conjunction with E2, transfers the ubiquitin to a lysine residue on the protein substrate by an isopeptide bond. The conjugated ubiquitin can be further modified in a similar fashion, which ultimately leads to polyubiquitination of the protein substrate and degradation by the 26S proteasome^9^. Substrate specificity in this process is governed by the E3 ligase. Therefore, many of the current targeted protein degradation technologies repurpose an E3 ligase to bind to the specific protein of interest (POI). One approach is to use small molecules such as PROTACs (PROteolysis-TArgeting Chimeras) or molecular “glues” to bridge an E3 ligase with the protein substrate^10,11^. However, a different small molecule would be needed for each ligase-substrate pair, limiting the applicability of PROTACs for studying protein function in basic research.

Tag-based methods offer a more generalized approach to degrading any POI to address basic biology questions as well as for therapeutic target validation. Two different types of tags are commonly used: destabilizing amino acid sequences, commonly referred to as degrons, are used to tag the POI and promote interaction with endogenous E3 ligases in the presence of drugs^12^. Alternatively, larger tags, such as EGFP, are used to tag the POI and to recruit a modified E3 ligase, whose natural substrate-binding domain has been replaced with a nanobody against EGFP^13–15^.

So far, most of these methods rely on mammalian E3 ligases, particularly Cereblon (CRBN) and von Hippel-Lindau (VHL), which may not be highly expressed or functional in all cell types or tissues^16,17^. Furthermore, it was recently reported that different mammalian E3 ligases show variable degradation efficiencies with different substrate proteins^18^. Besides limitations of the E3 ligase utilized, another challenge in applying the existing degron-based methods to target an endogenous protein is the large size of most of the tags utilized (up to hundreds of amino acids^19–22)^, potentially interfering with the normal POI function and causing inefficient knock-in at the endogenous POI loci. The smallest tags reported so far are derived from IKZF3 (I3D) and SALL4 (S4D) and are 25 and 28 amino acids long, respectively^23,24^. Recently, Ludwicki and colleagues reported that replacement of the substrate recognition domain of a bacterial E3 ligase, IpaH9.8, with nanobodies against EGFP, could mediate more efficient degradation of EGFP-tagged proteins in mammalian cells when compared to commonly used mammalian E3 ligases^25^. We reasoned that a combination of a modified bacterial E3 ligase and smaller tags could offer a simplified approach to targeting endogenous proteins for degradation. We tested two small tags: a 15-amino-acid ALFA tag^26^ and a 11-amino-acid HiBiT tag^27^. Each of these tags have ready-to-use interacting partners, with a high affinity nanobody for the former (named NbALFA) and the LgBiT fragment for the latter. We show that when these interacting partners are fused to the catalytic domain of the bacterial E3 ligase IpaH9.8, their interaction with the ALFA or HiBiT tags mediates efficient degradation of tagged proteins in a reversible manner. We also show that in contrast to commonly used mammalian E3 ligases, <u>a</u>rtificial <u>b</u>acterial <u>E</u>3 ligases (ABELs) display high degradation efficiency with less variation among a range of different substrates tested. Finally, we show that when the substrate recognition domain of the IpaH9.8 is replaced with a high-affinity nanobody against human α-synuclein, it mediates efficient degradation of endogenous α-synuclein in cultured neurons as well as in humanized transgenic mice.

Our work demonstrates the flexibility and efficiency of tag-dependent and independent ABEL-based platforms in mediating efficient degradation of various substrates *in vitro* and *in vivo*, with great potential for drug target discovery and validation.

## RESULTS

### An artificial bacterial E3 ligase can be directed to specific substrates using small tags

To test whether small tags and a modified bacterial E3 ligase (IpaH9.8) could degrade a POI (Fig. 1a), we utilized the 14-15 amino acid long “ALFA” tag, for which a high affinity nanobody (NbALFA) is available^26^. We first derived human HEK293T stable cell pools expressing a fusion protein consisting of mCherry, ALFA tag and human α-synuclein (Fig. S1a). We then transiently transfected these cells with a plasmid encoding an artificial bacterial E3 ligase (ABEL), in which the substrate binding domain LRR of IpaH9.8 is replaced with NbALFA. To facilitate analysis of transfected cells, a nuclear-localized tagBFP fluorescent protein is co-expressed with ABEL. After 24 hours, the cells were collected for flow cytometry (Fig. S1b). The median fluorescence intensity (MFI) level of mCherry in tagBFP-positive (tagBFP^pos^) cells were quantified and normalized to cells transfected with a plasmid that only encodes tagBFP. Interaction between the ALFA tag and its nanobody NbALFA brings the catalytic domain of ABEL in close proximity to the ALFA-tagged fusion protein, triggering its degradation (Fig. 1a). Indeed, more than 80% reduction of mCherry signal was observed in tagBFP^pos^ cells transfected with NbALFA-ABEL. NbALFA-ABEL mutants preventing interaction with ALFA tag (NbALFA*ABEL) or harboring a C337A mutation abolishing its E3 enzyme activity (NbALFA-dABEL) failed to reduce the mCherry signal, indicating that recruitment of E3 ligase activity is required for substrate degradation (Fig. 1b, c). Experiments with the ALFA tag at the N-terminus had similar results (Fig. S2). Together, these data show that a combination of a small tag and an ABEL can mediate efficient degradation of a POI. We name this approach <u>de</u>gradation by <u>s</u>mall tag <u>ABEL</u> or “DESTABEL”.

**Figure 1:**
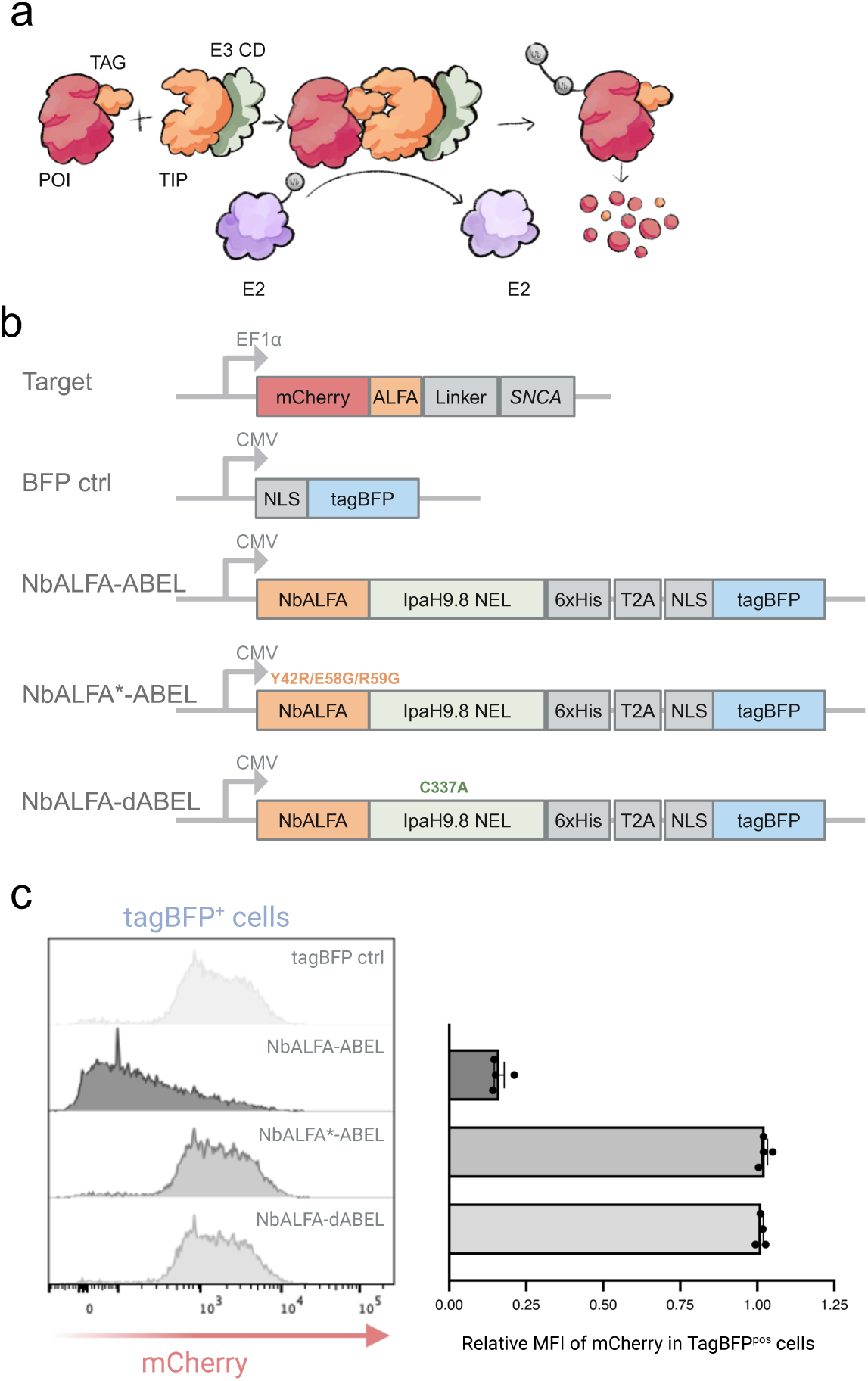
Efficient degradation of ALFA tag-containing fusion protein by an artificial bacterial E3 ligase. (a) Principle of degradation by small tag and artificial bacterial E3 ligase (DESTABEL). E3 CD, E3 ligase catalytic domain; POI, protein of interest; TIP, tag-interacting partner. (b) Diagram of plasmid constructs. The NbALFA of NbALFA*ABEL harbors three point mutations (Y424, E58G, R59G) that disrupt the interaction with the ALFA tag whereas NbALFA-dABEL loses catalytic activity due to a point mutation (C337A). (c) Flow cytometry data showing the loss of mCherry signal in BFPpos cells transfected with NbALFA-ABEL plasmids but not the control plasmids. The relative median fluorescence intensity (MFI) of mCherry in tagBFPpos cells is normalized to tagBFPpos cells transfected with a plasmid that only encodes tagBFP. N=4 biological replicates. Shown is mean and SEM.

### Reversible temporal control of protein degradation with DESTABEL

A major advantage to modulating gene function at the protein level is the potential for both temporal control and reversibility. We therefore next introduced a Doxycycline (Dox)-inducible NbALFA-based ABEL expression cassette into HEK293 cells stably expressing the mCherry-ALFA-human α-synuclein fusion protein (Fig. 2a). Using increasing concentrations of Dox, we first tested the dosage-response of fusion protein degradation. Dox induced expression of ABEL (as detected by tagBFP) and reduced mCherry fluorescence (Fig. 2b, e) in a dose-dependent manner. To test reversibility of degradation of the fusion protein, we first induced degradation of mCherry-ALFAtag-α-synuclein with Dox, then removed Dox to stop the production of ABEL. Within 2-3 days of Dox removal, cells regained mCherry fluorescence indicating the reversibility of protein degradation in our system (Fig. 2c, d, and f). Together, these data show that degradation of the fusion protein is tightly controlled temporally by Dox in a reversible manner.

**Figure 2:**
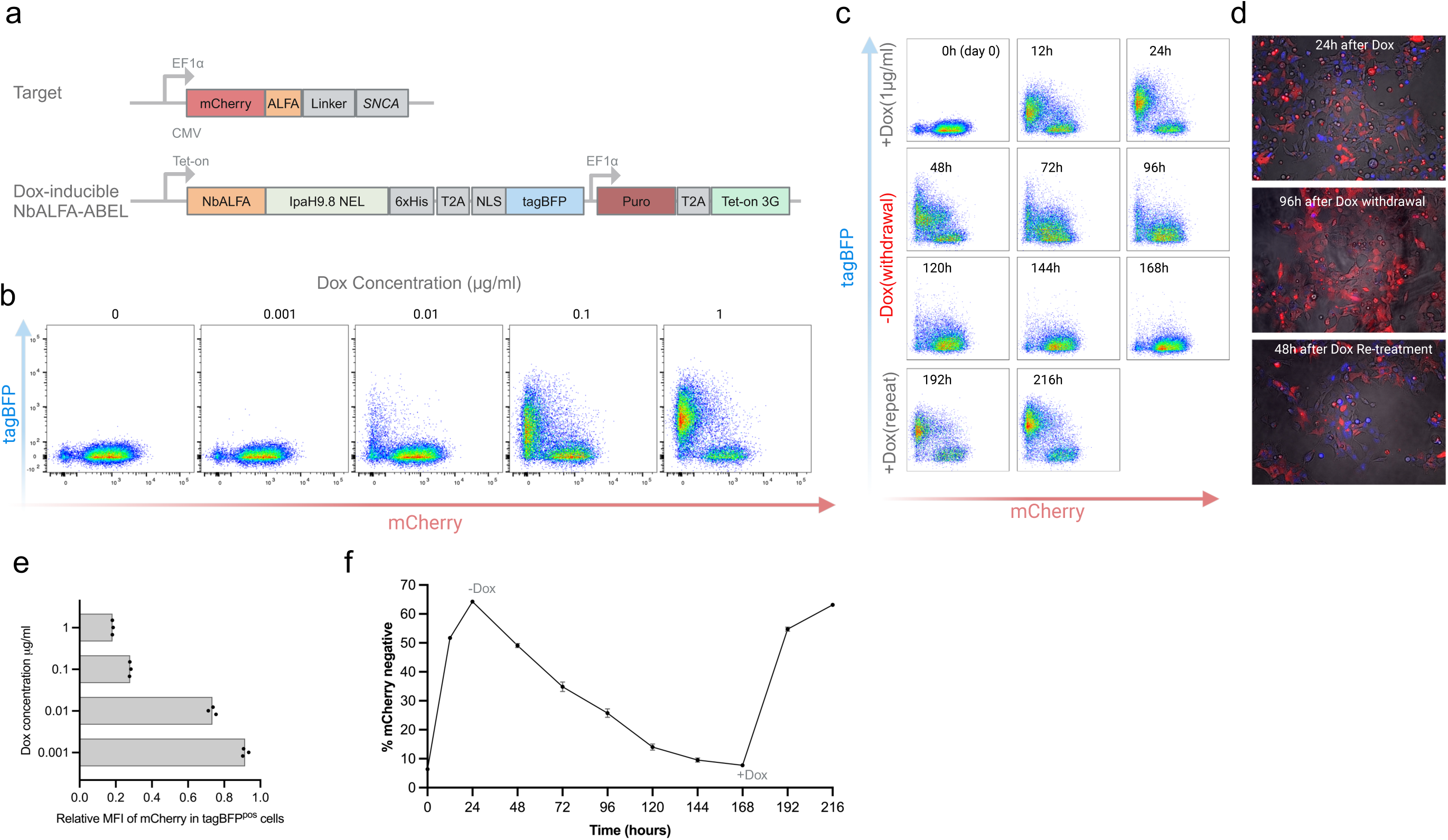
Dox-dependent reversible degradation of target protein. (a) Diagram of constructs used in HEK293 cells: constitutively expressed target protein mCherry-ALFAtag-synuclein and a Dox-inducible NbALFA-ABEL expression cassette; (b) Representative pseudo-colored dot plots from flow cytometry showing Dox-dependent degradation of the mCherry-ALFAtag-synuclein fusion protein. (c) Representative flow cytometry dot plots showing the time-course of Dox-induced degradation of mCherry. (d) Representative images at different time points showing the non-overlapping pattern between tagBFP (degrader) and mCherry (target protein). Please note that the cells are not clonal and not all cells respond to Dox induction. (e) Quantification of data shown in “b”. Scatter dot plots (n=3 biological replicates) of relative median fluorescent intensity (MFI) of mCherry in tagBFPpos cells with increasing concentration of Dox. Shown is mean and SEM. (f) Quantification of data shown in “c”. N=3 biological replicates and data shown is mean and SEM.

### DESTABEL can degrade endogenous substrates

Having shown that DESTABEL can be employed to efficiently degrade exogenous proteins, we next tested whether we could degrade an endogenous protein by tagging its corresponding genomic locus. We first established a polyclonal pool of mouse MC38 cells containing a stably integrated Dox-inducible ABEL expression cassette and then used these as parental cells for subsequent tagging of *Cd274*, encoding PD-L1, with ALFA tag (Fig. 3a, b). We used a CRISPR/Cas9-mediated HDR-based knock-in strategy to tag both alleles of *Cd274* at its C terminus (Fig. 3a). The small size of the ALFA tag (encoded by 42 bp) made it possible to use a single-stranded short oligo donor as the recombination template (Fig. 3a). Cells nucleofected with a gRNA/Cas9 RNP complex as well as the oligo donor were clonally-expanded to obtain homozygous edited cells. Dox-treatment of multiple clones showed approximately 80% reduction of PD-L1 protein (Fig. 3c-d and Fig. S3). Together, these data show that STABEL is a powerful tool for modulating endogenous protein level.

**Figure 3:**
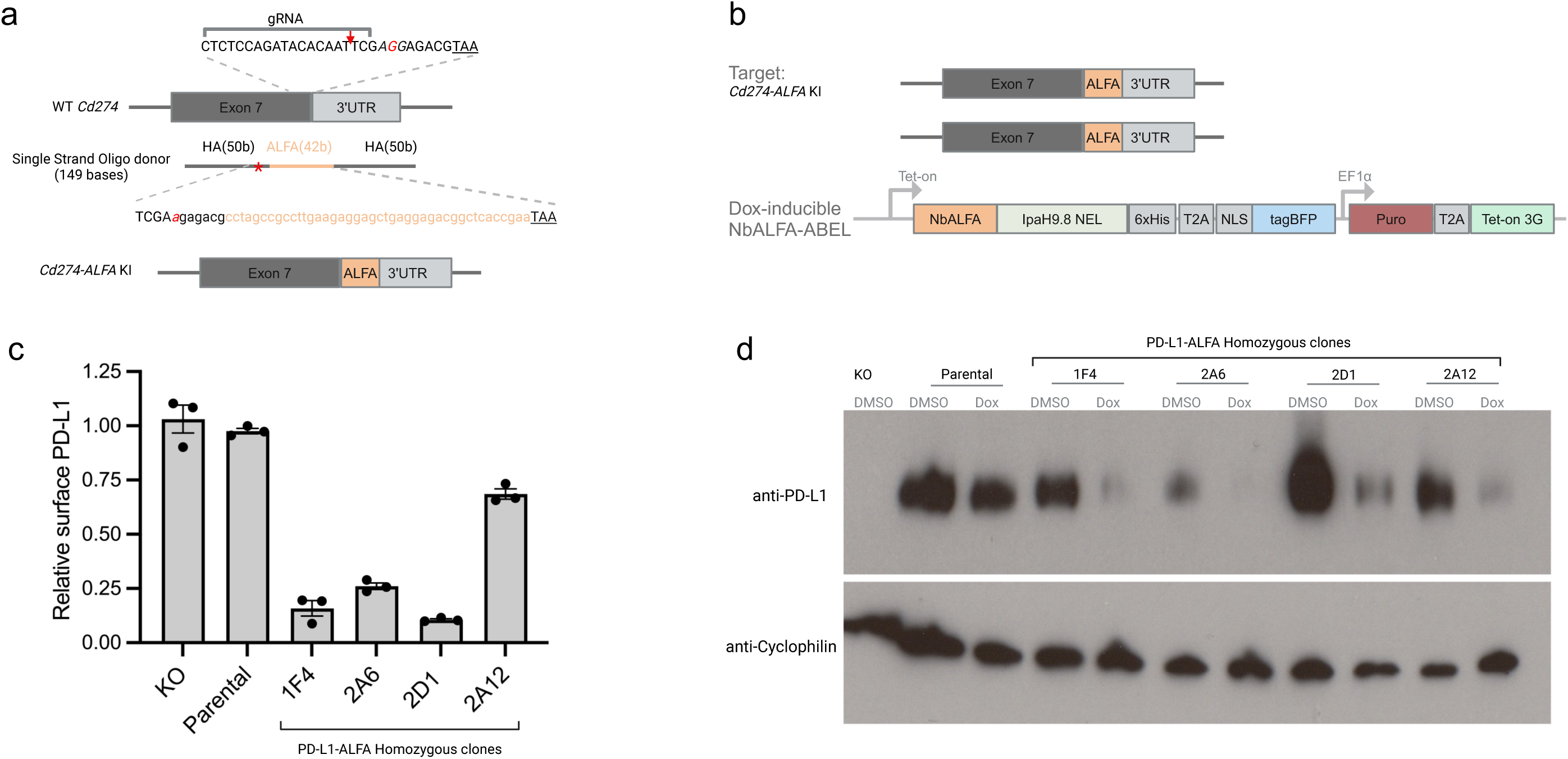
Tagging and Dox-controlled degradation of endogenous type I membrane protein PD-L1 in mouse MC38 cells with DESTABEL. (a) Strategy to tag the endogenous alleles of *Cd274* in mouse MC38 cells. Stop codon is underlined; PAM sequence is italicized; the cut site of the gRNA is denoted with the red arrow; the ALFA tag-coding sequence in the oligo donor is indicated. To prevent re-cutting by Cas9 after homologous-recombination, a single silent G to A mutation is introduced into the PAM sequence in the donor oligo (colored red in the sequence and denoted with red-asterisk in the diagram). (b) Diagram showing a homozygous clone containing a Dox-inducible NbALFA-based ABEL cassette as a transgene and two copies of ALFA-tagged endogenous *Cd274*. (c) Quantification of flow cytometry analysis of surface level PD-L1 following Dox-induced expression of ABEL in four homozygous clones. Surface PD-L1 level post Dox treatment is normalized to the same clone treated with DMSO. N=3 biological replicates. Shown is mean and SEM. (d) Western blot confirmation of Dox-induced degradation of PD-L1 in homozygous clones.

### HiBiT tag can be used for both targeted protein degradation and quantification

The LgBiT and HiBiT are split N-terminal and C-terminal fragments from nanoluciferase, respectively, which spontaneously interact to reconstitute luciferase activity. The 11 amino-acid NanoLuc-derived HiBiT has been used extensively to tag endogenous proteins for quantification purposes. HiBiT’s high affinity for the complementary LgBiT fragment allows for reconstitution of nanoluciferase activity in live cells, cell lysates and HiBiT-blotting buffer^27^. We reasoned that we could use the strong interactions between HiBiT and LgBiT to direct HiBiT-tagged proteins for LgBiT-based ABEL degradation (Fig. 4a). Since it was unclear whether fusion of the ABEL catalytic domain might affect the interaction between LgBiT and HiBiT, we tested different constructs with the LgBiT at either the N-terminus or C-terminus of the fusion protein. We made stable HEK293T cell pools expressing an mCherry-ALFA-HiBiT fusion protein and then transfected these cells with plasmids encoding LgBiT-based ABELs of various designs (Fig. 4b). We found that while all versions of LgBiT-ABELs can degrade the fusion protein irrespective of HiBiT’s position in the substrate, the N-terminus LgBiT-ABEL showed the highest degradation activity (Fig. 4c) and was comparable to ALFA-mediated degradation. Relative mCherry MFI in the presence of dABEL was in some instances above the level observed in the parental cells, possibly due to stabilization of mCherry in the presence of the LgBiT unit. Next, we investigated whether the same HiBiT tag can be used for both mediating and detection of protein degradation (Fig. 4d). Cells transfected with either NbALFA and LgBiT-based ABELs showed significant reduction of the protein substrate using LgBiT-based HiBiT blotting for detection, consistent with the anti-ALFA Western blot results (Fig. 4d). The denaturation and SDS-PAGE steps are sufficient to destroy the strong interactions between the HiBiT tag and the LgBiT-based ABELs, as shown in cells transfected with the LgBiT-based dABEL (note the weak band marked with red star, which is the residual complex between the fusion protein and LgBiT-dABEL).

**Figure 4:**
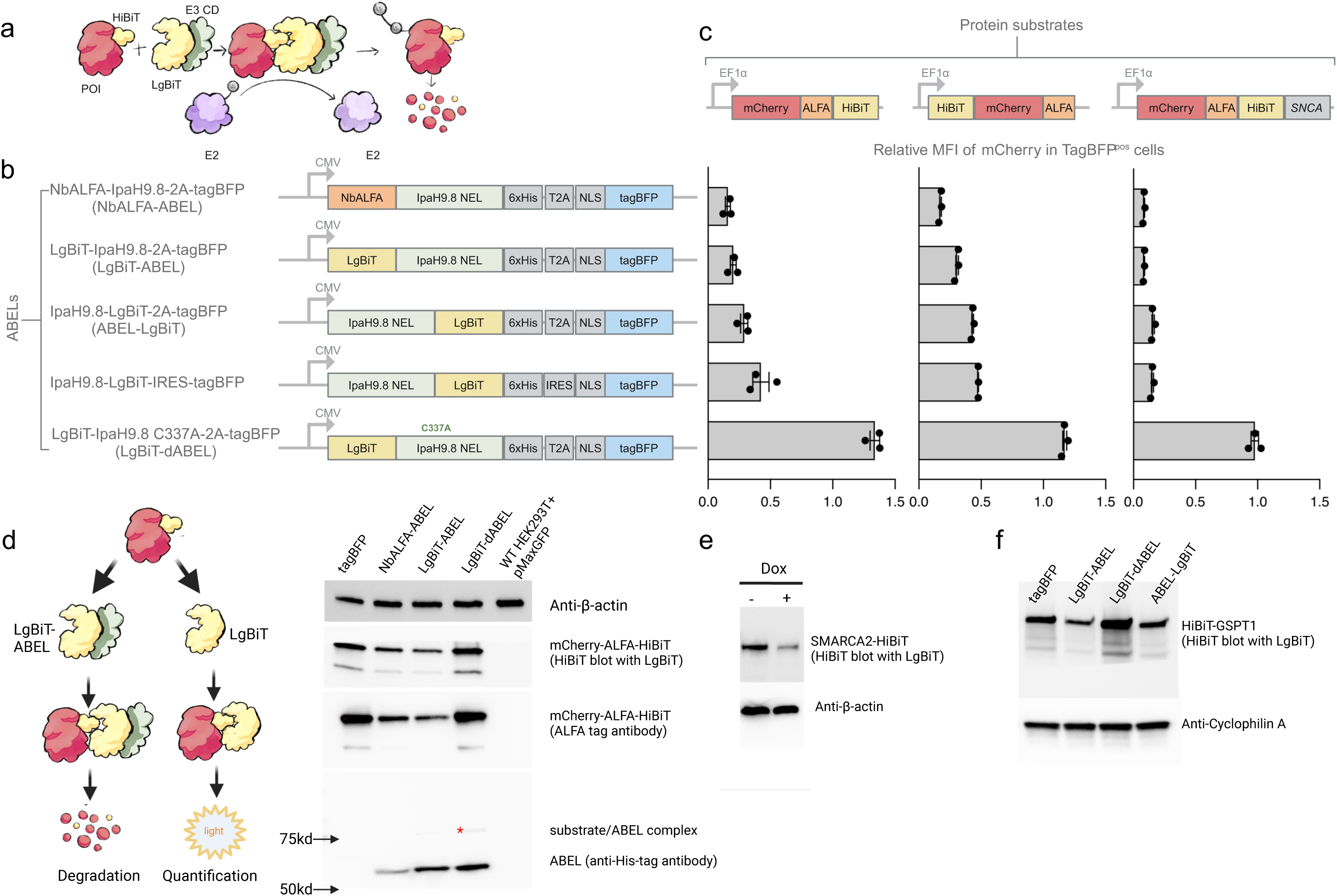
HiBiT tag mediates degradation of target protein with LgBiT-based ABEL. (a) Principle of HiBiT/LgBiT-based degradation. (b) Diagram of LgBiT-based ABELs. (c) Quantification of target protein degradation by FACS. N=3 biological replicates. Shown is mean and SEM. (d) The HiBiT tag can be used both for degradation of the fusion protein mCherryALFA-HiBiT and quantification of the protein degradation with HiBiT blotting. Please note the 2nd panel is performed with LgBiT-based HiBiT blotting whereas the rest with antibody-based conventional Western blot. Red star denotes residual fusion protein/LgBiT-ABEL complex, which is only visible with dead LgBiT-ABEL and not wild-type LgBiT-ABEL. Wildtype HEK293T cells transfected with a control plasmid (pMaxGFP) was included as a background control. (e) Western blot showing degradation of endogenously tagged SMARCA2 protein in HeLa cell lines upon Dox-induced expression of the LgBiT-based ABEL. (f) Western blot showing degradation of HiBiT-tagged endogenous protein GSPT1. Please note that the presence of a constitutively expressed LgBiT in the cell does not seem to abolish degradation. The cells were not sorted post transfection.

We next tested whether LgBiT-ABEL could also be used to degrade HiBiT-tagged endogenous protein. We introduced a Dox-inducible LgBiT-ABEL-expressing plasmid into human HeLa cells where one allele of *SMARCA2* was tagged with the HiBiT tag. Upon Dox treatment, efficient degradation of SMARCA2 was observed when compared to control cells treated with DMSO, demonstrating that LgBiT-ABEL can mediate efficient degradation of HiBiT-tagged endogenous protein (Fig. 4e). This was further confirmed with HiBiT-tagged GSPT1 protein in HEK293 cells (Fig. 4f). Note that presence of the LgBiT protein from the expression of a stably integrated plasmid, which was introduced for easy quantification of HiBiT-tagged GSPT1 in cell lysate, does not prevent LgBiT-ABEL mediated degradation. Similar to what was observed with overexpressed exogenous protein (Fig. 4c), position of the HiBiT tag in the endogenous protein at either the N-terminus (HiBiT-GSPT1) or C-terminus (SMARCA2-HiBiT) does not affect degradation (Fig. 4e, f).

### IpaH9.8-based ABEL consistently outperforms artificial mammalian E3 ligases (AMELs)

It has been reported that AMEL efficiency is substrate dependent^18^. We decided to investigate whether ABEL might behave differently and generated six different HEK293T stable cell pools expressing different protein substrates, all containing a fluorescent protein (tdTomato, mCherry or EGFP) and the ALFA tag (Fig. 5a). We then transfected each of the stable cell pools with six different plasmids encoding either IpaH9.8-based ABEL or SPOP, KLHL6, FBXL15, CRBN, or VHL-based AMELs (Fig. 5b). The location of NbALFA in the fusion proteins is determined by the location of the substrate binding domain of each AMEL. While SPOP-AMEL, KLHL6-AMEL and VHL-AMEL could efficiently degrade the TDP43 Q331K-EGFP-ALFAtag fusion protein, degradation efficiency of the other fusion proteins was variable. In fact, most of the fusion proteins could not be efficiently degraded by the majority of the AMELs, although these mammalian E3 ligases are either frequently used (SPOP, VHL, CRBN) or have been described as insensitive to substrate change^18^. In contrast, in our system the IpaH9.8-based ABEL consistently shows efficient degradation of all substrates tested, regardless of subcellular localization (Fig. 5c), consistent with the literature^25^. These data suggest that the IpaH9.8-based ABEL is a more robust tool for degradation of most POIs in mammalian cells.

**Figure 5:**
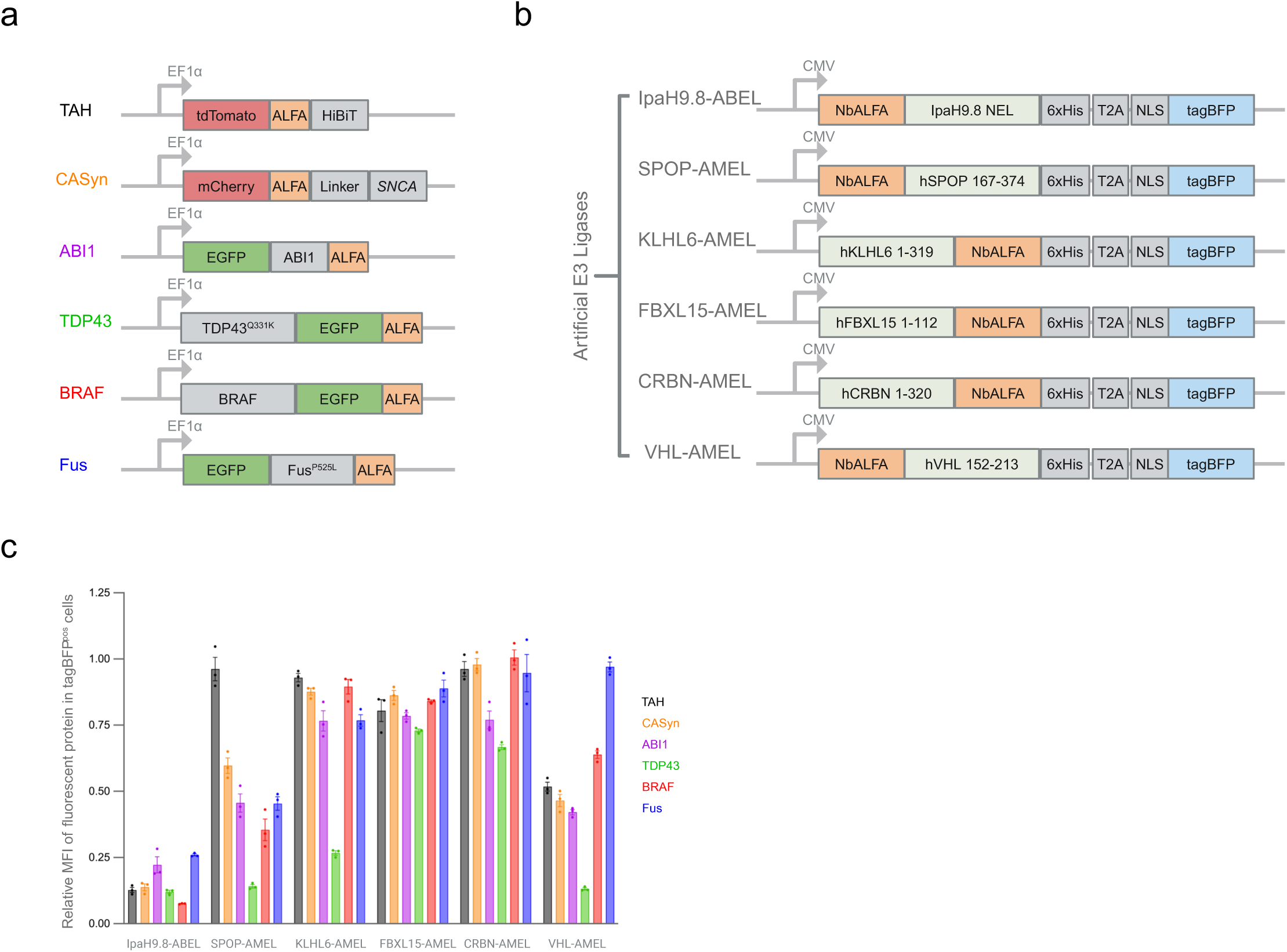
Consistent high degradation efficiency of different substrates by IpaH9.8-based ABELs. (a) Six different target fusion protein substrates. (b) Bacterial IpaH9.8-based ABEL was compared to that of five different mammalian E3 ligase-based AMELs. (c) Quantification of MFI relative to tagBFP alone. N=3 biological replicates. Shown is mean and SEM.

### ABEL can degrade un-tagged proteins via a target-specific nanobody moiety

Artificial E3 ligases consisting of a protein-specific nanobody and the ligase catalytic domain have been shown to be able to degrade multiple proteins involved in cancer, including KRAS and PCNA^28–30^. The consistent degradation efficiency of ABEL observed so far encouraged us to test whether similar strategies could be used to degrade proteins expressed in the brain. Having already shown that fusion proteins containing α-synuclein (encoded by *SNCA*) can be degraded with a tag-specific nanobody, we decided to focus on α-synuclein for which multiple high affinity nanobodies exist^31–34^ (Fig. 6a-c). HEK293 cells stably expressing a fusion protein between mCherry, ALFA tag and human α-synuclein were transiently transfected with plasmids encoding various ABELs, consisting of a nanobody moiety against either mCherry (LaM4)^35^ or human α-synuclein (NbSyn87, NbVH14 and NbSyn2) (Fig. 6d-e). ABELs consisting of the mCherry nanobody LaM4 (LaM4-ABEL) and α-synuclein nanobody NbSyn87 (NbSyn87-ABEL) could mediate an 80% reduction of the fusion protein, while others were ineffective (Fig. 6d). It is not clear why NbVH14 and NbSyn2 were not mediating degradation; we speculate that NbSyn2’s relatively lower affinity for α-synuclein monomers (kd ∼264nM for NbSyn2 at 37°C vs. ∼42nM for NbSyn87) might be part of the explanation (similar data are not available for NbVH14.) Alternatively, intrinsic differences may exist among different nanobodies that determine their ability to recruit ABEL for degradation as it has been reported that NbSyn87 and NbSyn2, although both bind to the C-terminal region, have different effects on the aggregation of α-synuclein *in vitro*^36^. As expected, degradation relies on the catalytic activity of the E3 ligase domain (Fig. 6d)^25^. We also observed that a short 7 amino acid linker (GSGSGSS) between NbSyn87 and ABEL completely abolishes the activity of NbSyn87ABEL (Fig. 6d), suggesting that the proximity between ABEL and the fusion protein substrate is critical. Western blot with either anti-mCherry or anti-synuclein antibody confirmed degradation of the fusion protein (Fig. 6e).

**Figure 6:**
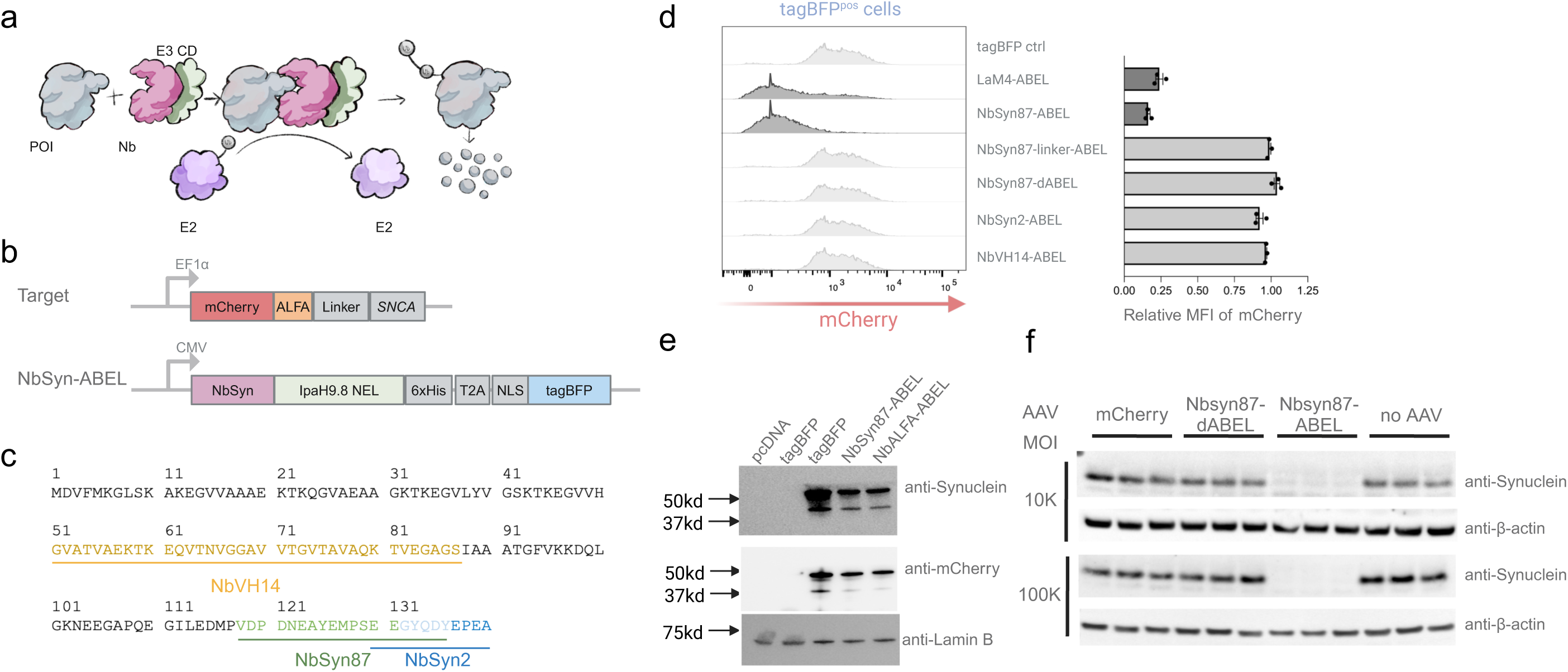
ABEL, consisting of a nanobody against α-synuclein (NbSyn87), mediates efficient degradation of human α-synuclein in cultured cells. (a) Diagram shows how an endogenous protein of interest can be degraded by an ABEL containing a target protein-specific nanobody. (b) Plasmids encoding an α-synuclein-containing fusion protein target and a αsynuclein nanobody-based ABEL, respectively. (c) Amino acid sequence of human α-synuclein (SNCA) and the reported binding epitopes of three different nanobodies. (d) Representative flow cytometry analysis of protein degradation activity mediated by ABELs with various Nb moieties (left). Scatter dot plot quantification of target protein degradation relative to cells transfected with tagBFP alone (right). N=3 biological replicates. Shown is mean and SEM. tagBFP ctrl: control construct without ABEL. LaM4: anti-mCherry Nb. (e) Representative western blot showing ABEL-mediated protein degradation. Lanes 1+2: wildtype HEK293 cells, lanes 3-5: HEK293 cells stably expressing the mCherry-ALFAtag-synuclein fusion protein. (f) Western blot showing NbSyn87-based ABEL-mediated degradation of human α-synuclein in cultured primary neurons derived from human *SNCA* BAC transgenic mice and infected with AAVs carrying ABEL fusion protein constructs.

We next examined whether wild-type α-synuclein could also be degraded by NbSyn87ABEL. We first derived stable HEK293T cells overexpressing human α-synuclein and transfected them with various ABELs. Consistent with what we observed in the flow cytometry experiments with mCherry-ALFA-α-synuclein, significant degradation of overexpressed α-synuclein could only be observed with NbSyn87-ABEL, but not NbSyn2-ABEL and VH14-ABEL (Fig. S4a). Similarly, cells transfected with NbSyn87-dABEL do not show any reduction of α-synuclein (Fig. S4a). Consistent with previously published IpaH9.8 data showing that degradation is mediated by the 26S proteasome^37^, treatment of cells with proteasome inhibitor MG132 blocked the reduction of α-synuclein (Fig. S4a). We also tested whether endogenously expressed α-synuclein could be degraded by NbSyn87-ABEL. Two human malignant melanoma-derived cell lines, MeWo and SK-Mel-28^38^, that both express α-synuclein, were transiently transfected with control tagBFP, NbSyn87-ABEL and NbSyn87-dABEL plasmids. Transfected cells were enriched by FACS and α-synuclein level was assessed by Western blot. Consistent with our previous observations, for both cell lines, those transfected with NbSyn87-ABEL, but not NbSyn87-dABEL, show reduction of α-synuclein (Fig. S4b).

To explore degradation of human α-synuclein in a physiologically relevant cell type, we next used primary cortical neurons from Bacterial Artificial Chromosome (BAC) transgenic mice expressing human *SNCA*. We infected these primary neurons with Adeno-Associated Viruses (AAVs) encoding mCherry, NbSyn87-ABEL or NbSyn87-dABEL (Fig. S5). Western blotting show titer-dependent reduction in endogenous human α-synuclein in a ligase activity-dependent manner (Fig. 6f). These data demonstrate that AAV-mediated delivery of NbSyn87-ABEL results in highly efficient degradation of human α-synuclein in cultured mouse primary neurons.

### ABEL-mediated degradation of α-synuclein *in vivo*

After demonstrating that NbSyn87-ABEL can mediate efficient degradation of human α-synuclein in cultured primary transgenic mouse neurons, we tested whether efficient degradation could also be achieved *in vivo.* The transgenic mice used in this study express wildtype human synuclein at a lower level than endogenous mouse synuclein. Hence there are no aggregates in this transgenic mouse model. We retro-orbitally injected PHP.eB-based AAVs encoding either mKate2, NbSyn87-ABEL or NbSyn87-dABEL into adult human BAC *SNCA* transgenic mice (Fig. 7a). We find that mKate2 expression is reduced when positioned after T2A and therefore likely under-reports degrader expression. 12 weeks (cohort 1) and 10 weeks (cohort 2) post AAV injection, animals were sacrificed and brain tissues were collected for analysis (Fig. 7b). ELISA quantification showed ∼43% reduction in human α-synuclein abundance in the NbSyn87-ABEL cohort when compared to mKate2 (about 11 μg/g brain tissue vs. about 6 μg/g brain tissue) (Fig. 7c). In contrast, NbSyn87-dABEL did not reduce human α-synuclein protein, indicating specific protein degradation dependent on ligase activity. This result was further confirmed with Western blot analysis from tissue lysates of brain cortex (Fig. S6). Immunostaining with a human α-synuclein-specific antibody confirms the reduction in the brains injected with NbSyn87-ABEL-expressing virus, but not in the controls (Fig. 7d). Quantification of α-synuclein expression in different brain regions suggests that significant degradation occurs in the neurons of cortex, hippocampus and hypothalamus regions where α-synuclein is uniformly expressed in these transgenic mice and a degradation trend was observed in the thalamus as well (Fig. 7e).

**Figure 7:**
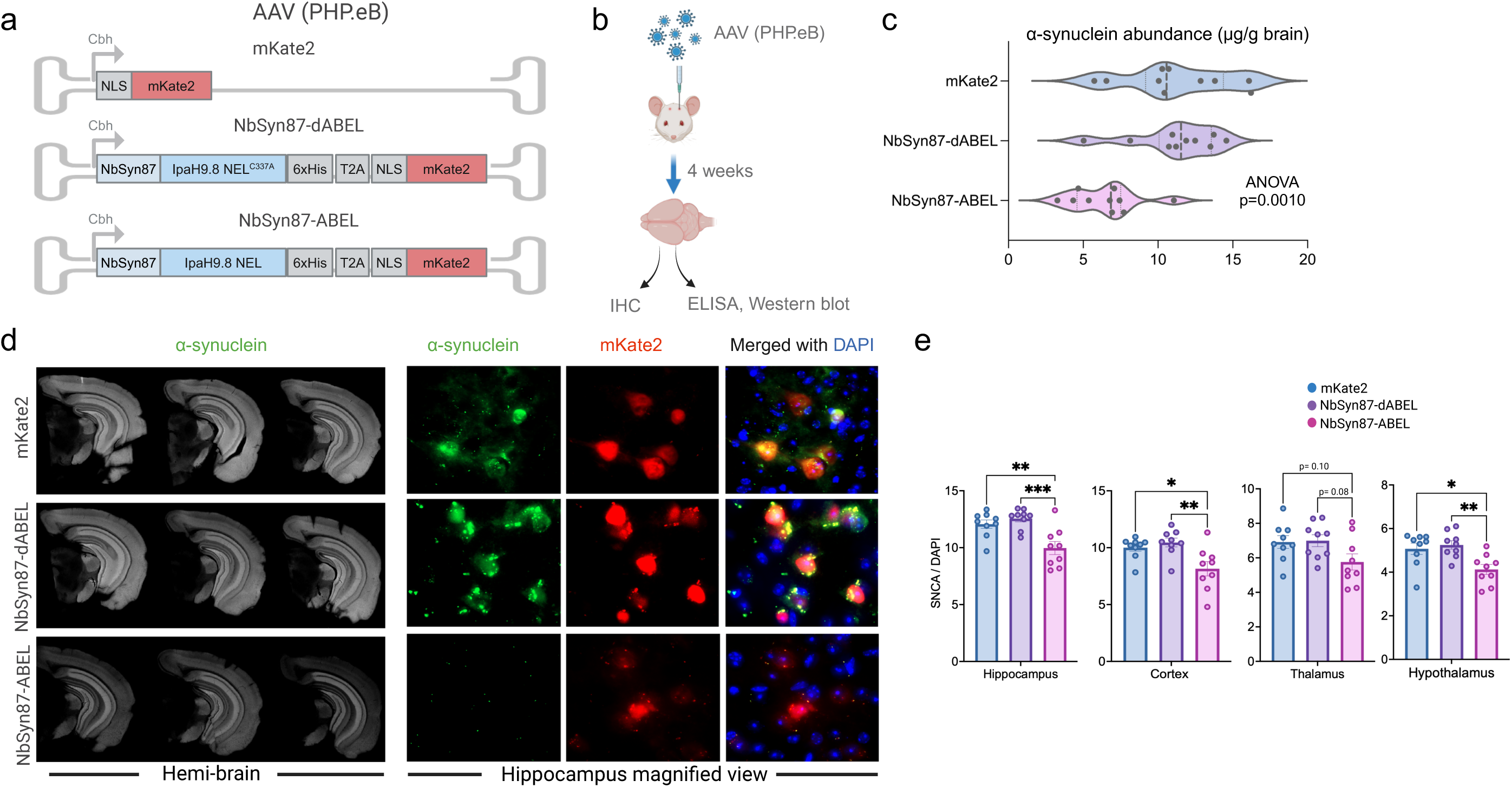
Reduction of human α-synuclein after PHP.eB-AAV-based delivery of NbSyn87-ABEL into the adult brain of transgenic mice. (a) Diagram of PHP.eB AAV viruses used to deliver NbSyn87-ABEL, a control version with dead E3 ligase catalytic domain (NbSyn87-dABEL), or mKate2 reporter only. (b) Experimental workflow. (C) Violin plots showing ELISA measurement of human α-synuclein level in brain lysates (n=10 mice in each group). Comparison was done using one-way ANOVA analysis. (d) Representative immunostaining with human α-synuclein-specific antibody and mKate2 fluorescence (e) Quantification of α-synuclein expression in different brain regions (n= 9 animals in each group, one-way ANOVA followed by Bonferroni’s multiple comparison test, p-value 0.05). Asterisks (*) denote pairwise statistical significance.

Together, these data demonstrate that AAV-based delivery of NbSyn87-ABEL can degrade human α-synuclein *in vivo*.

## DISCUSSION

Modulating the function of endogenous genes at the protein level offers unique advantages when compared to manipulations at the genomic and mRNA level. Existing technologies for protein modulation often involve the use of large degrons, which makes tagging of endogenous genes cumbersome. Here we show that small tags, relatively easily and efficiently knocked into the endogenous loci with CRISPR-based homology-directed recombination, coupled with an exogenously expressed artificial bacterial E3 ligase consisting of a tag-interacting partner and the catalytic domain of a bacterial E3 ligase, can simplify this process. We also show, using a dox-inducible expression system, that we are able to reversibly induce the degradation of a protein of interest. Reversibility of protein degradation, in contrast to the irreversible complete loss achieved with DNA-level knockout, may better mimic drug effects and offer the opportunity to model different dosing schemes in pre-clinical studies.

In this study, we tested two small tags, the ALFA tag and the HiBiT tag. It is worth noting that knock-in of such tags into an endogenous gene offers extra benefits in addition to controlled degradation. For example, with various commercially-available reagents, the presence of the ALFA tag may enable easy purification or immunoprecipitation of the protein of interest and fluorescent super-resolution microscopy imaging of its expression in cells^26^. Similarly, we show that the HiBiT tag can both mediate degradation of the tagged protein and be used to quantify degradation efficiency with a simple HiBiT blot. The latter is particularly useful for detecting proteins with no available antibodies. We propose that other small tags, such as the 12aa SPOT tag, which is derived and optimized from β-catenin and has a known high affinity nanobody^39^, may be used for similar purposes as the ALFA tag. However, some small tags with identical epitopes found in endogenous proteins such as the 10aa 5B9^40^ should be used with caution due to the potential for off-target degradation.

Different from most degron-based systems, we took advantage of the bacterial E3 ligase IpaH9.8, which has previously been shown to have superior degradation activities over commonly used mammalian E3 ligases^25^. Furthermore, it was reported that mammalian E3 ligases may show substrate dependent degradation efficiency^18^. Our data confirmed these observations and show that IpaH9.8 seems devoid of such degradation efficiency variation, at least for the substrates tested here. We also note that a linker between the nanobody and the catalytic domain of the IpaH9.8 is not only unnecessary, but sometimes could be detrimental, suggesting that the proximity between the substrate protein and the ABEL is critical for the degradation. While this manuscript was in preparation, Mercer and colleagues reported the evolution of a 36 amino acid zinc finger-based degron (SD40)^41^, which seems to be devoid of non-specific (off-target) degradation. However, the SD40 platform is dependent on endogenous CRBN E3 ligase for targeted protein degradation. *CRBN* gene expression varies greatly among different tumor cell lines and could even be completely absent^16^. Furthermore, most mammalian E3 ligases or exogenously expressed plant auxin receptor TIR1 require the co-operative action of multiple endogenous proteins to form a functional complex^9^. In contrast, to our knowledge, there is no evidence suggesting that the bacterial E3 ligase IpaH9.8 depends on endogenous mammalian proteins to function properly. Therefore, our system, relying on an exogenous ABEL, would be less susceptible to the mutation/expression of endogenous E3 ligase components.

Alternatively, our system may be used orthogonally with SD40 to modulate the degradation of two different endogenous proteins. Orthogonal degradation might also be achievable with other bacterial E3 ligases^42^.

Neuronal proteins such as ɑ-synuclein, amyloid-beta and tau are expressed in the brain and are susceptible to aggregation. The aggregated forms of these proteins are pathological hallmarks of several neurodegenerative diseases^43,44^, including Parkinson’s and Alzheimer’s diseases. As a proof of concept, we show here that a nanobody-based ABEL can be used to degrade ɑ-synuclein *in vivo*, which in-turn might limit the formation of aggregated proteins. Out of scope for this study, one potential application of the system could be to examine the effect of protein degradation on protein aggregation kinetics. Based on other studies and approaches used to reduce the specific disease-causing proteins in the brain^45,46^, we speculate that this strategy could be very effective in attenuating disease progression in the brain.

While an artificial bacterial E3 ligase is an unlikely therapeutic candidate, we envision that our DESTABEL platform will greatly enable the process of tagging endogenous genes and modulate their cognate protein function in a temporally controlled manner in both cell culture and *in vivo*, which might be particularly useful for target discovery and validation of therapeutic targets before significant resources are devoted to the development of PROTACs or molecular glues.

## METHODS

### Primers, gRNAs, Plasmids and AAVs

All PCR primers and gRNAs (crRNAs and tracrRNA) were ordered from Integrated DNA Technologies (IDT). PCR analyses were carried out with AccuPrime Taq DNA polymerase HiFi (ThermoFisher Scientific). Full length gRNAs were reconstituted by annealing crRNAs to tracrRNA according to manufacturer’s protocol and their activity was assessed by transfecting cells followed by a T7E1 assay (IDT). All plasmids were synthesized, cloned and purified by Genscript Inc. AAV1 and PHP.eB-AAV were packaged and prepared by Virovek Inc. Plasmid sequences are available upon request.

### Cell Culture

HEK293T, MC38, HeLa, MeWo and SK-MEL-28 cells were obtained from Genentech’s internal quality-controlled cell line repository (CellCentral). HEK293T, MC38, HeLa and SK-Mel-28 cells were cultured in DMEM supplemented with 10% FBS and 2mM GlutaMAX; MeWo cells were cultured in RPMI-1640 supplemented with 10% FBS and 2mM GlutaMAX. HEK293T cells were either transfected with Lipofectamine 2000 (Thermo Fisher Scientific Inc) or nucleofected with solution F (Lonza Inc) according to manufacturer’s recommendations; HeLa cells were nucleofected with solution E (Lonza Inc) according to manufacturer’s recommendations; MeWo and SK-Mel-28 cells were transfected with Lipofectamine 2000 (Thermo Fisher Scientific Inc).

To establish stable cell pools expressing fluorescent fusion proteins, HEK293T cells were co-transfected with a PiggyBac-configured plasmid containing the coding cassette for the fusion protein and a plasmid encoding the PiggyBac transposase at a 3:1 mass ratio. The cells were cultured/passaged for 10 days before FACS-sorted to enrich for mCherry^pos^ cells. To introduce Dox-inducible ABEL degrader plasmids, a similar approach was utilized except that puromycin selection instead of FACS was employed to enrich for the desired cells.

### Tagging of endogenous *Cd274* alleles in mouse MC38 cells

A MC38 cell line that constitutively expresses RFP-Luciferase fusion protein was cultured in DMEM supplemented with 10% FBS and 2mM GlutaMAX. The cells were first electroporated with a PiggyBac-configured plasmid containing the Dox-inducible NbALFA-ABEL degrader cassette and a plasmid encoding the PiggyBac transposase at a 3:1 mass ratio via nucleofection (solution E with program CM138) according to manufacturer’s instructions (Lonza Inc.), followed by puromycin selection (5μg/ml) three days post nucleofection. 2x10^5^ puromycin-resistant cells were then nucleofected with 5μl RNP complex (2μl TruCut Cas9 protein V2 at 5μg/μl; 3μl sgRNA at 50μM and 1μl oligo donor at 100μM). An aliquot of the nucleofected cells were analyzed 3 days later for homologous recombination efficiency and the rest of the cells were sorted into 96-well plates to derive single cell clones. About 3 weeks later, wells with single clones were screened by PCR with primers flanking the insertion region. Due to the small size difference between the PCR products of the wild type allele and the knock-in allele, 10% Novex TBE PAGE gel (EC62752BOX, ThermoFisher Scientific) was used for DNA electrophoresis. Homozygous clones were confirmed by Sanger sequencing and expanded for further analysis.

### Flow cytometry and FACS

HEK293T cells stably expressing fluorescent fusion protein were nucleofected with plasmids encoding various ABELs and analyzed by flow cytometry 24 hours later. PD-L1 wild-type, knockout and ALFA-tagged cells were cultured in the presence of 20 ng/ml interferon-γ (R&D Systems 485-MI), treated with DMSO or 1 µg/ml Dox for 24 hours before flow cytometry. Data were collected on a BD FACSCelesta cell analyzer with FACSDiva software (BD Inc) and analyzed with FlowJo (BD Inc.). Sorting of transfected MeWo and SK-Mel-28 cells was performed on a BD FACSAria Fusion cell sorter.

### ELISA

Left cerebrum of each brain was weighed and lysed in RIPA buffer supplemented with Complete protease inhibitor (Roche Diagnostics Inc) at a ratio of 1 ml buffer per 100 mg tissue with Qiagen TissueLyser II (Qiagen Inc). Tissue lysate was centrifuged at 12,500 rpm for 15 minutes and supernatant (generally with a total protein concentration around 7 μg/μl) was serially diluted by factors of 10 to reach a 1,000x dilution, which was further serially diluted by factors of 2 to 2,000x, 4,000x, 8,000x and 16,000x dilution. The last four dilutions were used for ELISA with Legend Max Human alpha-synuclein (colormetric) ELISA kit (Biolegend Inc.) according to manufacturer’s protocol. Plates were read with SpectralMax i3x (Molecular Devices Inc.) and data analyzed with Graphpad Prism 10.

### Western blot

Cultured or sorted cells were lysed in 1xLaemmli buffer supplemented with 2-mercaptoethanol. Supernatants of brain lysates prepared as above at a total protein concentration around 7 μg/μl were mixed with equal volume of water and two times volume of 2xLaemmli buffer supplemented with 2-mercaptoethanol and boiled for 5-10 minutes. Samples were then loaded onto and separated with 4-20% Mini-Protean TGX Precast SDS-PAGE gel (Bio-Rad Inc), transferred to nitrocellulose membrane, blocked and detected with primary antibody followed by HRP-conjugated secondary antibody. To increase the detection sensitivity of human α-synuclein, the nitrocellulose membrane was fixed in 0.4% paraformaldehyde for 30 minutes before being blocked and detected with primary antibody. The following primary antibodies were used: polyclonal rabbit anti-synuclein antibody (Novus biologicals NBP2-15365, 1:2,000 dilution; for Western blot in Fig. 6e); human α-synuclein-specific mouse antibody (31C2, developed in-house^47^; 1:10,000 dilution, for the rest of the α-synuclein Western blots); polyclonal rabbit anti-mCherry antibody (Novus biologicals NBP2-25157, 1:250 dilution); monoclonal mouse anti-β-actin antibody AC-15 (Sigma-Aldrich A5441, 1:10, 000); rabbit monoclonal anti-lamin B1 antibody (Invitrogen #702972, 1: 1,000 dilution); polyclonal rabbit anti-tubulin antibody (Cell Signaling Technology #2144, 1:1,000 dilution). HRP-conjugated goat anti-rabbit (ThermoFisher Scientific #32460, 1:20,000 dilution) and goat anti-mouse (ThermoFisher Scientific #31430; 1:20,000 dilution).

Primary mouse cortical neurons were washed with PBS and lysed with RIPA lysis buffer (Thermo Fisher, 89901) containing benzonase nuclease (Millipore Sigma, E1014) and protease/phosphatase inhibitor cocktail (Thermo Fisher, 78440). Cells were centrifuged at 15,000x *g* for 5 minutes, and supernatants were saved. Lysates were heated to 95°C with NuPAGE LDS sample buffer (Invitrogen, NP0007) for 5 minutes and loaded onto NuPAGE 4-12% Bis-Tris Midi Gels (Invitrogen, WG1403BOX) and run at 140V for approximately 1.5 hrs. Samples were transferred to nitrocellulose membranes (BioRad, 1704159), blocked with 5% BSA in PBST for one hour at room temperature, probed with primary antibody overnight at 4°C, washed with PBST and probed with secondary antibody for one hour at room temperature. Antibodies used: Syn211 (Thermo-Fisher Scientific, #32-8100); Beta-actin (Cell Signaling Technology, #3700).

### HiBiT blotting

HiBiT blotting was performed following manufacturer’s protocol (Promega, N2410).

### Primary mouse cortical neuron cultures

Cortices from both male and female day 15 embryos (E15.5) were dissected, stripped of meninges, washed 3x with cold HBSS (Invitrogen, 14170-112), and incubated for 10 min at 37°C in HBSS supplemented with 0.25% trypsin (Invitrogen, 15090-046) and DNase I (Roche, 104159). Tissue was washed 3x with HBSS and triturated in plating media containing DNase I (Neurobasal Medium (Thermo Fisher Scientific, #21103-049), 20% heat-inactivated horse serum (Thermo Fisher Scientific, #26050-163 088), 25 mM sucrose, and 0.25% Gibco GlutaMAX (Thermo Fisher Scientific, #35050-061). Dissociated cells were centrifuged at 125xg for 5 min at 4°C, resuspended in plating medium, and plated in poly-L-lysine-coated (Millipore Sigma, P1274) plates. After 24 hrs, the plating medium was replaced with NBActiv4 (Brain Bits, NB4500). Cells were maintained at 37°C with 5% CO_2_ and the medium was renewed using 50% exchange every 3-4 days. At day 7, AAV (AAV1 serotype) was added to the primary neuron culture, at an MOI of either 10K or 100K. 10 days later, cells were collected for immunoblotting.

### Mouse husbandry, AAV injection and tissue collection

All animal procedures and protocols were approved by the Institutional Animal Care and Use Committee at Genentech Inc. All animals were bred and maintained in a pathogen-free animal facility under standard animal room conditions. Two cohorts of human SNCA-wildtype BAC transgenic mice (C57BL/6N-Tg(SNCA)129Mjff/J, <u>JAX 018442</u>), each with 15 individuals of both sexes, were divided into three groups. Each mouse was retro-orbitally injected with 100 μl of AAVs at a titer of 1 x 10^13^ vc/ml. The AAVs were synthesized (Virovek, CA) in AAV2^PHP.eB^ serotype, carrying one of three constructs: pX52-Cbh-mKate2, pX52-Cbh-NbSyn87-IpaH9.8-T2A-mKate2, or pX52-Cbh-NbSyn87-IpaH9.8^C337A^-T2A-mKate2. The mice were sacrificed 12 and 10 weeks post-injection for the first and second cohorts, respectively. After cardiac perfusion with cold PBS, the brains were extracted. Each brain was dissected, with one hemisphere being flash-frozen on dry ice for later biochemical assays and the other hemisphere fixed in 4% paraformaldehyde in PBS for 24 hours, then transferred into PBS for later tissue sectioning and immunohistochemistry.

### Total Synuclein Immunohistochemistry

Fixed hemi-brains were sent to NeuroScience Associates Inc, where they were incubated with 20% glycerol and 2% Dimethyl Sulfoxide (DMSO) overnight to prevent any freeze artifacts. The hemi-brains were embedded and prepared for coronal sectioning within a gelatin matrix utilizing MultiBrain® Technology (NeuroScience Associates, Knoxville, TN). The hemi-brains were sectioned coronally at a thickness of 30 μm via a microtome. Next, the sections were subjected to immunohistochemistry. After a series of washes with PBS and PBST, sections were blocked for one hour using a room temperature solution of 5% Donkey Serum (DS). Post-blocking, the sections were incubated overnight at 4°C with the primary antibody (monoclonal anti human SNCA 31C2 (7.5 mg/ml) developed internally at Genentech^47^ at 1:2,000 dilution in 0.5% DS. The following day, sections were washed with TBST and then treated with a donkey anti-mouse secondary antibody (Thermo Fisher Scientific, #A-31571, 1:1,000) for two hours at room temperature. Following another pair of PBST washes, sections were mounted with a gelatin solution and air-dried. Lastly, a DAPI-containing ProLongTM Gold Antifade mounting medium (Thermo Fisher Scientific, #P36931) was applied, the specimen was overlaid with a cover slip, and left to dry overnight.

### Tissue analysis and Quantification

Slides were imaged using an Olympus VS200 slide scanner microscope at 4x magnification. DAPI was visualized at 360 nm, SNCA at 647 nm and mKate2 at 580 nm, respectively. The cortex, hippocampus, thalamus, and hypothalamus were identified using QuPath for whole slide analysis. Mean intensity of SNCA was divided by area and normalized to DAPI intensity for each region, then averaged for each animal. Individual cell images in brain tissues were captured using a Zeiss Apotome 3 at 40x magnification.

### Statistical analysis

All data were analyzed with Microsoft Excel or Graphpad Prism 10. One way ANOVA analyses were performed post-hoc with Microsoft Excel and Graphpad Prism 10.

### Figure production

Figure elements were produced in Microsoft Excel, Flowjo, and Graphpad Prism 10. Final figures were assembled using BioRender.com.

## ACKNOWLEDGEMENTS AND DISCLAIMER

We would like to thank the Genentech FACS lab for assistance with cell sorting, Kimberly Stark for coordinating NSA sample transfer, Neuroscience Associates Inc. (Knoxville, TN, USA) for providing the tissue processing infrastructure to conduct this study, Drs. Bruno Alicke and Xiaofen Ye for help with experiments, Dr. Honglin Chen for assistance with AAVs, and Drs. Peter Dijkgraaf, Robert Yauch, Anthony Partridge, Enrico Cannavò, and Tao Sun, for helpful discussions. We would also like to thank Miss Autumn Ruoran Liu for help with preparation of some of the diagrams. All authors are current or former employees of Genentech Inc., a member of the Roche Group.

---

**Figure S1:**
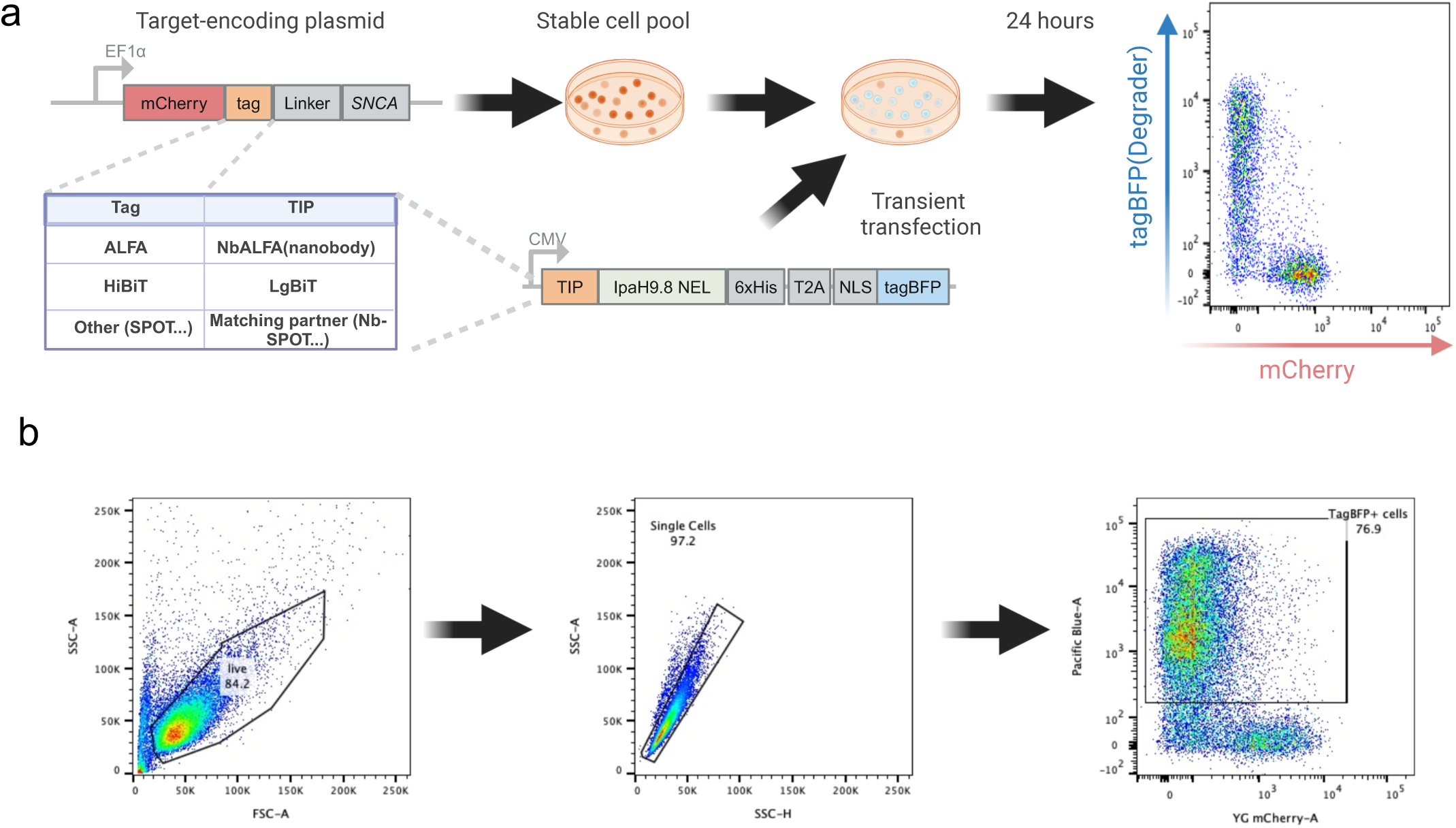
Schematic illustration of experimental procedures employed to test degradation of a POI. (a) General structure of constructs encoding target and degrader as well as experimental approach (b) A representative example of flow cytometry gating strategy.

**Figure S2:**
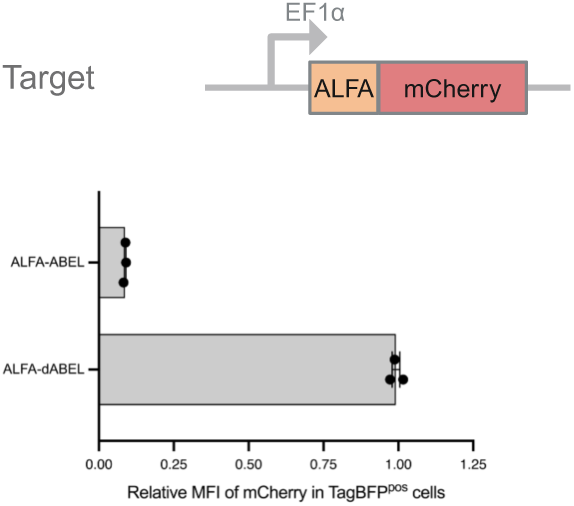
Efficient degradation of N-terminally ALFA-tagged fusion protein by NbALFA-ABEL. Scatter dot plot quantification of mCherry degradation relative to cells transfected with tagBFP alone. N=3 biological replicates. Shown is mean and SEM.

**Figure S3:**
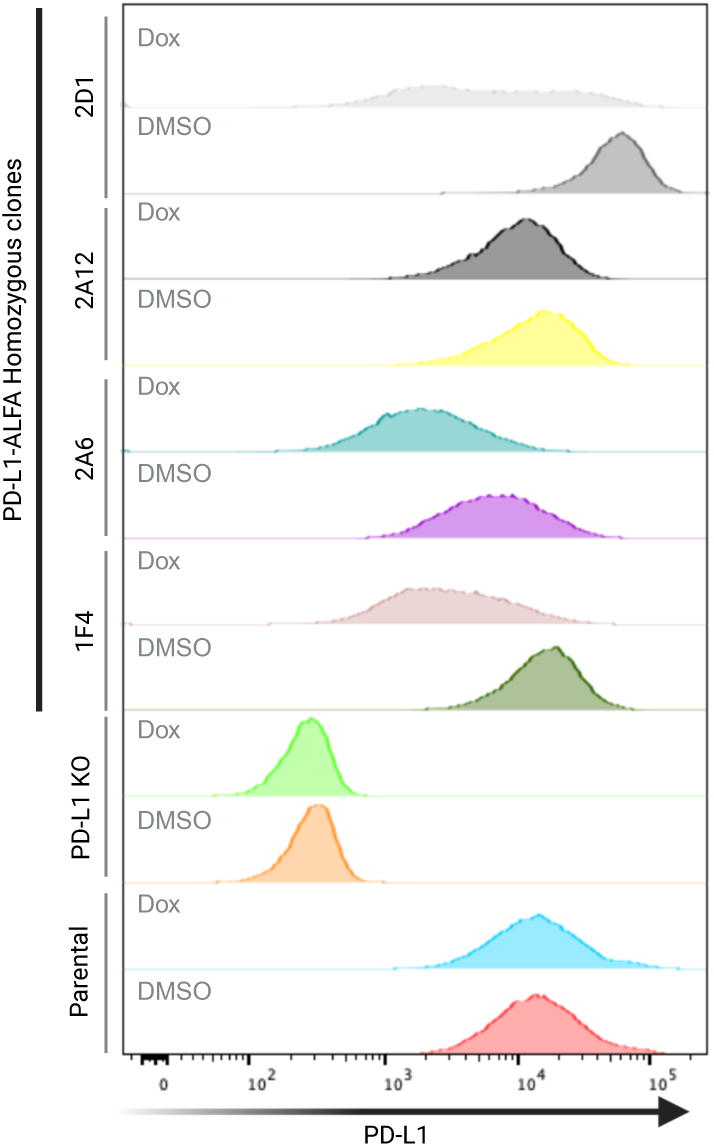
Efficient degradation of ALFA-tagged endogenous PD-L1. Representative flow cytometry data showing the level of surface PD-L1 in control (DMSO) and Dox-treated MC38 cells.

**Figure S4:**
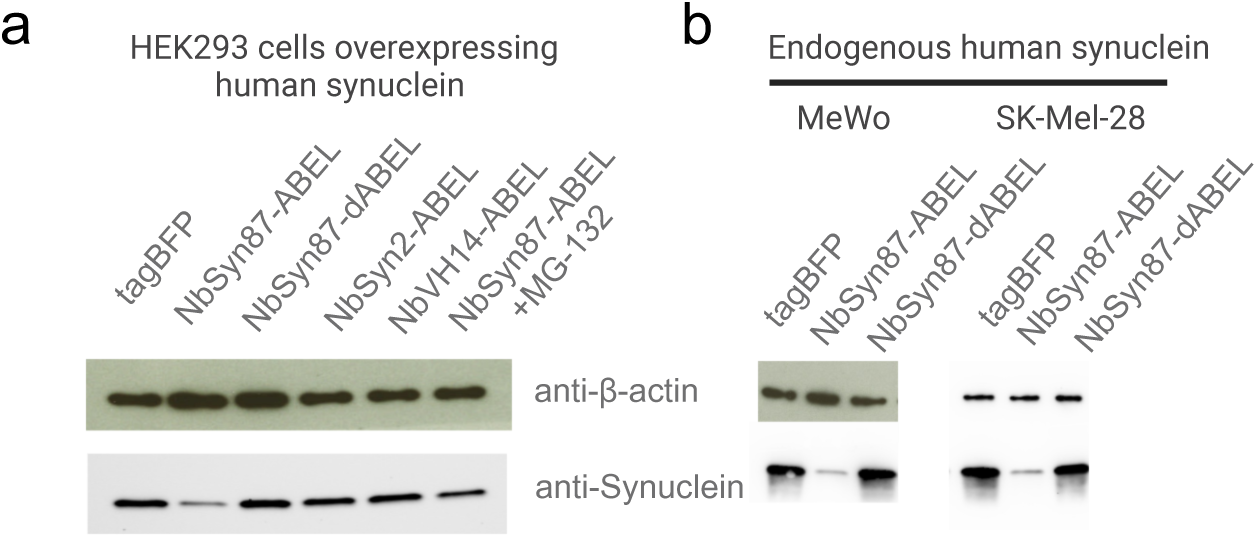
Efficient degradation of human α-synuclein. (a) Western blot showing degradation of human α-synuclein when overexpressed in HEK293 cells. (b) Western blot showing degradation of endogenous human α-synuclein in MeWo and SK-Mel-28 cells.

**Figure S5:**
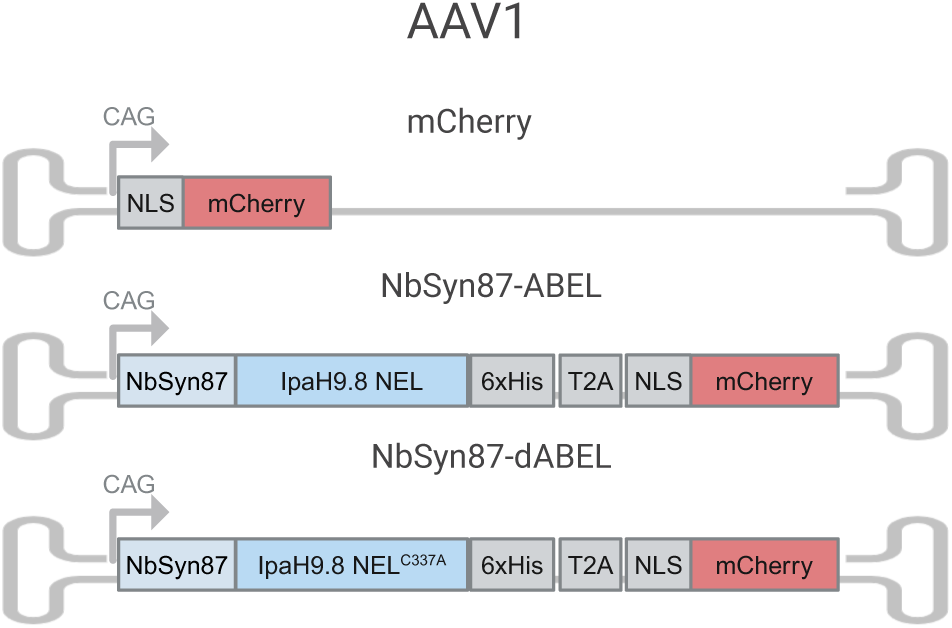
Diagram of AAV1 viruses used to infect cultured primary neurons. CAG promoter and mCherry reporter were used in these experiments.

**Figure S6:**
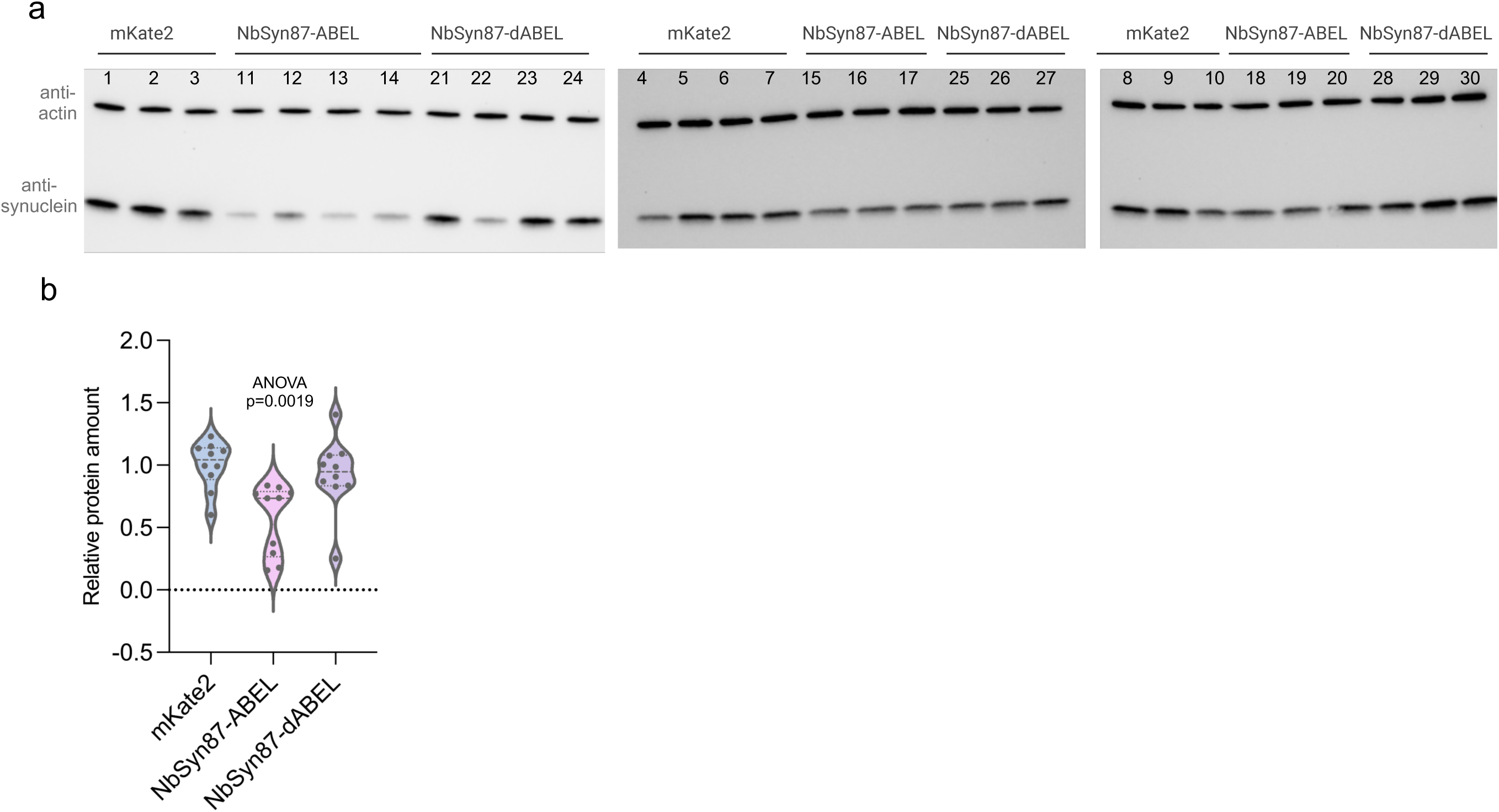
Human α-synuclein degradation in transgenic mouse brains injected with PHP.eB virus encoding NbSyn87-ABEL. (a) Western blot detection of human α-synuclein in mouse brain lysates with human α-synuclein-specific antibody; anti-β-actin staining serves as loading control; (b)Violin plots showing quantification of data in “a” , normalized to β-actin loading control. N=10 animals in each group.

**Table S1:**
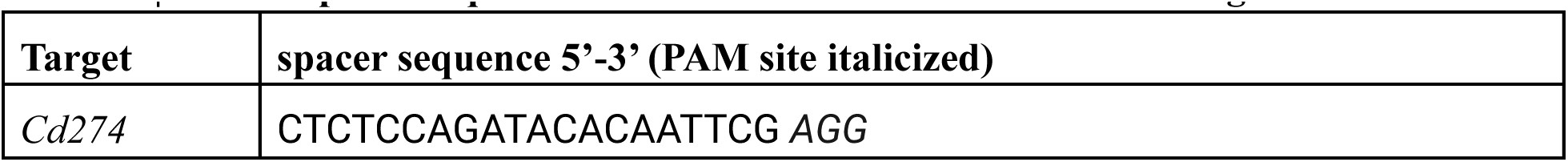
crRNA spacer sequence used to facilitate insertion of ALFA tag.

**Table S2:**
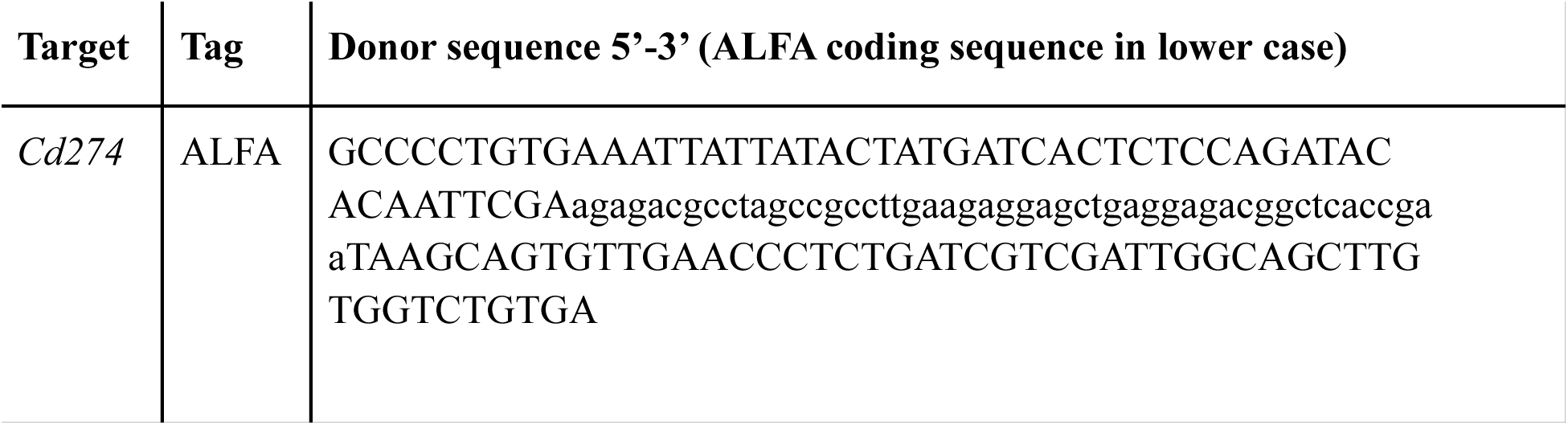
ssDNA donor template for ALFA knock-in.

**Table S3:**
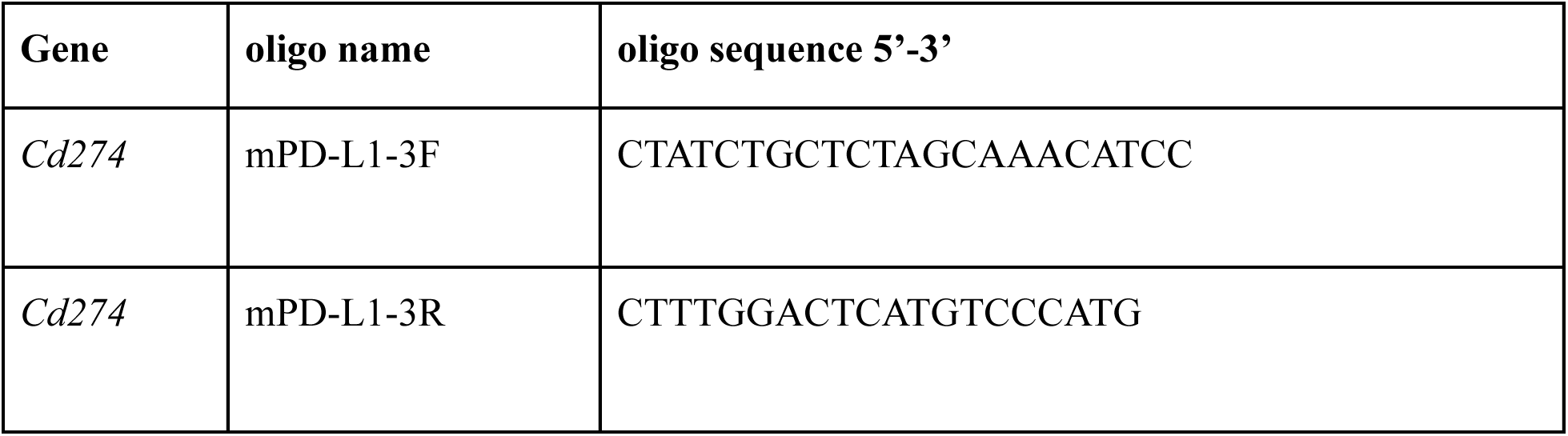
PCR primers used for Tag insertion validation.

